# Mrr1 regulation of methylglyoxal catabolism and methylglyoxal-induced fluconazole resistance in *Candida lusitaniae*

**DOI:** 10.1101/2020.05.18.101840

**Authors:** Amy R. Biermann, Elora G. Demers, Deborah A. Hogan

**Affiliations:** Department of Microbiology and Immunology, Geisel School of Medicine at Dartmouth, Hanover, NH 03755

## Abstract

In *Candida* species, the transcription factor Mrr1 regulates azole resistance genes in addition to the expression of a suite of other genes including known and putative methylglyoxal reductases. Methylglyoxal (MG) is a toxic metabolic byproduct that is significantly elevated in certain disease states that frequently accompany candidiasis, including diabetes, kidney failure, sepsis, and inflammation. Through the genetic analysis of *Candida lusitaniae* (syn. *Clavispora lusitaniae*) strains with different Mrr1 variants with high and low basal activity, we showed that Mrr1 regulates basal and/or induced expression of two highly similar MG reductases, *MGD1* and *MGD2*, and that both participate in MG detoxification and growth on MG as a sole carbon source. We found that exogenous MG increases Mrr1-dependent expression of *MGD1* and *MGD2* in *C. lusitaniae* suggesting that Mrr1 is part of the natural response to MG. MG also induced expression of *MDR1*, which encodes a major facilitator protein involved in fluconazole resistance, in a partially Mrr1-dependent manner. MG significantly improved growth of *C. lusitaniae* in the presence of fluconazole and strains with hyperactive Mrr1 variants showed greater increases in growth in the presence of fluconazole by MG. In addition to the effects of exogenous MG, we found knocking out *GLO1*, which encodes another MG detoxification enzyme, led to increased fluconazole resistance in *C. lusitaniae*. Analysis of isolates other *Candida* species found heterogeneity in MG resistance and MG stimulation of growth in the presence of fluconazole. Given the frequent presence of MG in human disease, we propose that induction of *MDR1* in response to MG is a novel contributor to *in vivo* resistance of azole antifungals in multiple *Candida* species.

**Author Summary:** In *Candida* species, constitutively active variants of the transcription factor Mrr1 confer resistance to fluconazole, a commonly used antifungal agent. However, the natural role of Mrr1 as well as how its activity is modulated *in vivo* remain poorly understood. Here, we have shown that, in the opportunistic pathogen *Candida lusitaniae*, Mrr1 regulates expression and induction of two enzymes that detoxify methylglyoxal, a toxic metabolic byproduct. Importantly, serum methylglyoxal is elevated in conditions that are also associated with increased risk of colonization and infection by *Candida* species, such as diabetes and kidney failure. We discovered that methylglyoxal causes increased expression of these two Mrr1-regulated detoxification enzymes as well as an efflux pump that causes fluconazole resistance. Likewise, methylglyoxal increased the ability of multiple *C. lusitaniae* strains to grow in the presence of fluconazole. Several other *Candida* strains that we tested also exhibited stimulation of growth on fluconazole by methylglyoxal. Given the physiological relevance of methylglyoxal in human disease, we posit that the induction of fluconazole resistance in response to methylglyoxal may contribute to treatment failure.

## Introduction

*Candida* species are among the most prominent fungal pathogens, with mortality rates for candidaemia ranging from 28 to 72% depending on geographic location (reviewed in (1)), and recent decades have seen a worldwide increase in the overall incidence of candidemia (2). Treatment failure of invasive fungal infections remains an important clinical issue (3) due to long-term complications, high mortality rates, and elevated healthcare costs. Perplexingly, treatment may fail even in cases where isolates from a patient have tested as susceptible to a certain antifungal *in vitro*. Therefore, it is clear that multiple cryptic factors which are not present *in vitro* may influence the outcome of antifungal therapy.

In *Candida* species, one mechanism of azole resistance is overexpression of the gene *MDR1* (4–7), which encodes an efflux pump. Overexpression of *MDR1* is usually caused by gain-of-function mutations in the gene encoding the zinc-cluster transcription factor Mrr1 (8–10). Many studies in *C. albicans* have focused on the relationship between Mrr1, *MDR1*, and fluconazole resistance (8, 10–13), but little is known about other genes that Mrr1 regulates. Thus, the natural role of Mrr1 is not well understood. By studying the biological functions of Mrr1-regulated genes, it is possible to gain insight into important questions such as the evolutionary purpose of Mrr1, drivers of selection for gain-of-function mutations in Mrr1, and other consequences of high Mrr1 activity aside from drug resistance. Independent studies in *C. albicans* (13–16), *C. parapsilosis* (17), and *C. lusitaniae* (7, 18) have revealed genes that appear coordinately upregulated in fluconazole-resistant isolates with gain-of-function mutations in *MRR1*.

Previously, we demonstrated a link between fluconazole (FLZ) resistance and specific single nucleotide polymorphisms (SNPs) in the *MRR1* locus (*CLUG_00542*) among twenty clinical *Candida* (*Clavispora*) *lusitaniae* isolates from a single patient with cystic fibrosis (7). We observed multiple *MRR1* alleles containing gain-of-function mutations which were directly related to FLZ resistance. The presence of *MRR1* alleles conferring high FLZ resistance among this population was unexpected, as the patient had no prior history of antifungal use. Thus, we became interested in other potential factors that could have selected for gain-of-function mutations in *MRR1*. An RNA sequencing (RNA-Seq) analysis comparing several isolates with high- or low-activity *MRR1* alleles indicated many genes which may be regulated by Mrr1 in these isolates (7). In accordance with aforementioned studies (13–17), two of the potential Mrr1 target genes encoded putative methylglyoxal reductases. Recently, another study in *C. lusitaniae* has also shown that a putative methylglyoxal reductase is regulated by Mrr1 in FLZ-resistant clinical isolates (18).

Methylglyoxal (MG) is a cytotoxic carbonyl compound formed spontaneously as a byproduct of multiple metabolic processes in all known organisms (Fig. 1). It is a highly reactive electrophile that irreversibly modifies proteins, lipids, and nucleic acids in a nonenzymatic reaction known as glycation, resulting in cellular damage and stress (19, 20). Serum levels of MG are elevated in patients with diabetes (21–23), sepsis (24), and uremia (25–28) relative to healthy controls. Additionally, MG is thought to be generated during inflammation as part of the neutrophil respiratory burst (29). In the yeast *Saccharomyces cerevisiae*, there is a positive correlation between rate of glycolysis and MG production (30), with approximately 0.3% of the total glycolytic carbon flux being converted to MG (31). MG may be detoxified through a glutathione-dependent glyoxalase system (32) or through an NADH- or NADPH-dependent MG reductase (33) (Fig. 1). Disruption of the glyoxalase pathway in *S*. *cerevisiae* leads to accumulation of intracellular MG (34) and renders cells highly sensitive to exogenous MG (35). The first MG reductase described in yeast was *S*. *cerevisiae* Gre2 (36); more recently, the *C*. *albicans* protein Grp2/Mgd1, which is homologous to Gre2, was found to have MG reductase activity (37). Homologs of *MGD1* and *GRE2* appear to be regulated by Mrr1 in many fluconazole-resistant *Candida* isolates across multiple species and families (7, 13–18). The *C. lusitaniae* genome encodes seven putative MG reductases (18), making these genes an interesting topic of study.

**Fig. 1.**
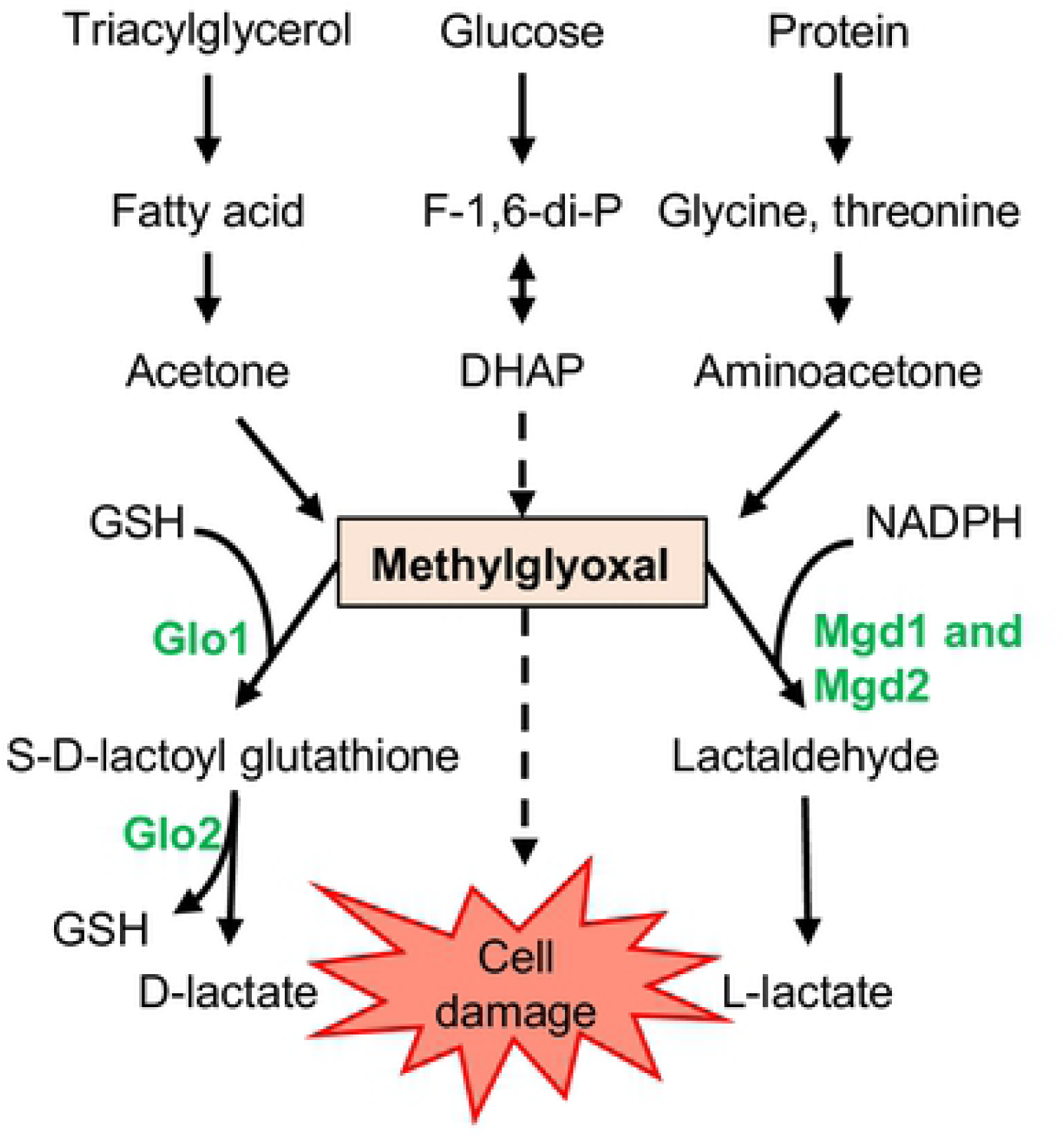
Methylglyoxal is a highly reactive, toxic byproduct that can form spontaneously during the catabolism of sugars, fatty acids, and proteins. It can be detoxified to D-lactate via the GSH-dependent glyoxalase system, consisting of Glo1 and Glo2, or to lactaldehyde through NAD(P)H-dependent methylglyoxal reductases such as Mgd1 and Mgd2 which are homologs of *C. albicans* Grp2. F-1,6-di-P, fructose 1,6-bisphosphate; DHAP, dihydroxyacetone phosphate; GSH, reduced glutathione. Solid arrows, enzymatic processes; dashed arrows, nonezymatic processes.

In the present study, we demonstrated that in *C. lusitaniae*, Mrr1 regulates basal and induced expression of *MGD1* (*CLUG_01281*) and induced expression of *MGD2* (*CLUG_04991*), both of which encode proteins important for detoxification and metabolism of MG. Deletion of one or both genes led to increased sensitivity to high concentrations of exogenous MG and decreased ability to use MG as a sole carbon source. In addition, we demonstrated that MG can induce Mrr1-dependent expression of *MGD1* and *MGD2*, as well as expression of *MDR1* in a partially Mrr1-dependent manner. MG increased growth in FLZ which was largely dependent on *MRR1* and *MDR1*. Additionally, we provided evidence that constitutively active Mrr1 leads to greater stimulation of growth in fluconazole compared to prematurely truncated Mrr1. A *glo1*∆ mutant was hypersensitive to MG but had higher FLZ resistance, consistent with the model that endogenous MG could activate *MDR1* expression. Finally, we showed that though MG sensitivity varies across *Candida* species, stimulation of azole resistance by MG is not exclusive to *C. lusitaniae*. To our knowledge, this is the first study to (i) experimentally examine the biological function of Mrr1-regulated genes other than the efflux pump genes *MDR1* and *FLU1*, (ii) establish a role for Mrr1 in a metabolic process, and (iii) demonstrate that MG, a metabolite associated with various human diseases, can increase resistance to a widely used antifungal agent.

## Results

### *Candida lusitaniae MGD1* and *MGD2* contribute to the detoxification and metabolism of MG

In our previous work, an RNA-seq analysis of clinical *C. lusitaniae* isolates showed that two genes with high sequence identity, *CLUG_01281* and *CLUG_04991*, were significantly more highly expressed in isolates with gain-of-function alleles of *MRR1* (7). The protein sequences encoded by *CLUG_01281* and *CLUG_04991* are 88% percent identical to each other and have 59% and 58% identity, respectively, to *C. albicans* Grp2 (Mgd1), a characterized MG reductase with homology and functional similarity to *S. cerevisiae* Gre2 (36, 37) (Fig. 2A). Based on homology to characterized MG reductases and experimental data shown below, from here forward *CLUG_01281* and *CLUG_04991* will be referred to as *MGD1* and *MGD2*, respectively. Sanglard and colleagues (18) reported an expansion of putative MG reductases and related aldehyde reductases in the *C. lusitaniae* genome. We further analyzed the relationships between putative MG reductases in select *Candida* species across various families, including *C. lusitaniae*, using FungiDB (38, 39). *MGD1* and *MGD2* are the two putative MG reductases most similar to *C. albicans GRP2/MGD1*. Other *Candida* species, including *Candida auris*, *Candida parapsilosis*, *Candida tropicalis* also have at least one set of similar putative MG reductases with at least 50% identity to *C. albicans* Grp2 (Fig. 2A). *C. glabrata* has a pair of related putative MG reductases that are homologous to *S. cerevisiae* Gre2 (Fig. 2A).

**Fig. 2.**
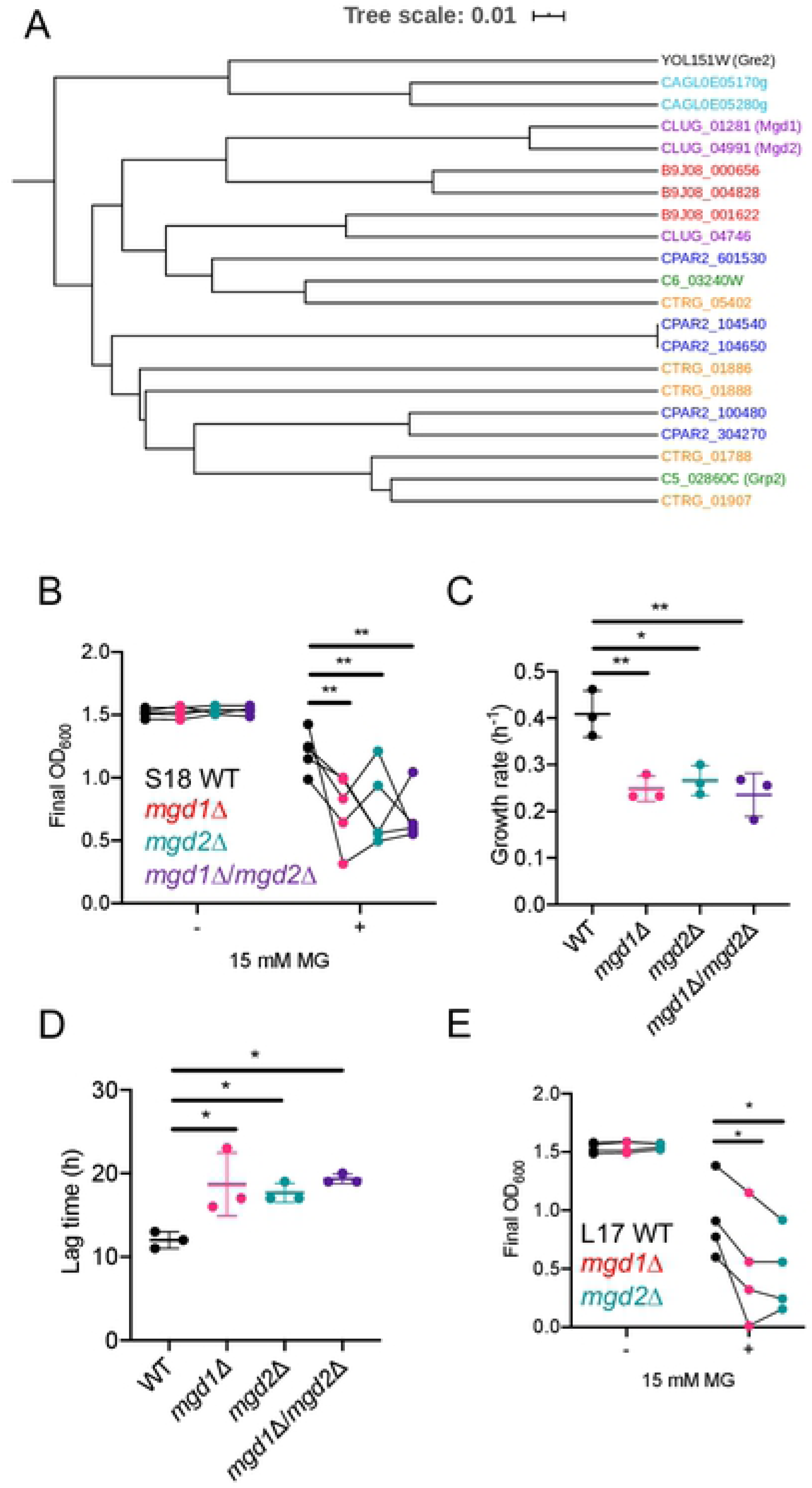
*MGD1* and *MGD2* are involved in MG detoxification. **(A)** Phylogeny of known and putative MG reductase amino acid sequences with homology to *C. albicans* Grp2, *S. cerevisiae* Gre2, and *C. lusitaniae* Mgd1 and Mgd2. *Candida* species is denoted by color: *Candida lusitaniae* (purple); *Candida auris* (red); *Candida tropicalis* (orange); *Candida parapsilosis* (blue); *Candida glabrata* (teal) and *Candida albicans* (green). **(B-D)** Growth of *C. lusitaniae* S18 WT (black), *mgd1∆* (magenta), *mgd2∆* (teal), and *mgd1∆*/*mgd2∆* (purple) strains in YPD with or without 15 mM MG in terms of final OD_600_ **(A)**, exponential growth rate **(B)**, and lag time **(C).** Data shown represent the mean ± SD from five independent experiments **(D)** Growth as final OD_600_ of strain L17 WT, *mgd1∆* and *mgd2∆* cultures in YPD with or without 15 mM MG. Data shown represent the mean ± SD from four independent experiments. A linear model was used for statistical evaluation of final OD_600_ in **(B)**. One-way ANOVA was used for statistical evaluation of data in **(C-E)**. * p < 0.05, ** p < 0.01. Connecting lines indicate each experiment.

To determine if *MGD1* and *MGD2* were involved in MG resistance and gain more insight into the respective roles of these two similar genes, we knocked out each gene independently and in combination in the clinical isolate S18, which contains a constitutively active Mrr1 variant (7). We grew the mutants and their parental isolate S18 (WT) in the presence or absence of 15 mM MG. MG caused an increase in lag phase and decreased growth rates and final yields when compared to control cultures. In the presence of MG, strains lacking *MGD1* and/or *MGD2* grew significantly worse than S18 WT (Fig. 2B-D and **S1C**). Specifically, the mutants had a 38 – 44% decrease in final yield (Fig. 2B), a 35 – 42% decrease in exponential growth rate (Fig. 2C), and a 48 – 61% increase in time spent in lag phase (Fig. 2D) relative to unaltered S18. None of the mutants displayed a growth defect in the control condition, suggesting that *MGD1* and *MGD2* are not essential in the absence of exogenous MG (Fig. 2B and **S1A**). Surprisingly, the double mutant did not exhibit a more severe phenotype than either single mutant indicating that the two enzymes are not redundant. To validate the phenotypes in a separate genetic background, we knocked out *MGD1* and *MGD2* in isolate L17 which also has constitutively active Mrr1. As with S18, knocking out *MGD1* or *MGD2* in L17 exacerbated the growth defects caused by MG (Fig. 2E and **Fig. S1D**). As expected, absence of *MGD1* and/or *MGD2* in either background did not affect FLZ minimum inhibitory concentrations (MICs) (Table 1).

**Table 1.** Fluconazole MIC of *C. lusitaniae* strains used in this paper

| Strain | FLZ MIC<br>(µg/mL) |
| --- | --- |
| S18 | 16 |
| S18 *mgd1*Δ | 16 |
| S18 *mgd2*Δ | 16 |
| S18 *mgd1*Δ/*mgd2*Δ | 16 |
| S18 *mrr1*Δ | 8 |
| S18 *cap1*Δ | 16 |
| S18 *mrr1*Δ/*cap1*Δ | 8 |
| S18 *mdr1*Δ | 2 |
| S18 *glo1*Δ | 16 |
| L17 | 16 |
| L17 *mgd1*Δ | 16 |
| L17 *mgd2*Δ | 16 |
| L17 *mrr1*Δ | 8 |
| L17 *cap1*Δ | 16 |
| L17 *mdr1*Δ | 4 |
| U04 | 32 |
| U04 *mrr1*Δ | 4-8 |
| U04 *mrr1*Δ + *MRR1*-<br>Y8 | 32 |
| U04 *mrr1*Δ + *MRR1*-<br>L1Q1* | 0.25 – 0.5 |
| U05 | 0.5 - 1 |
| L14 | 0.5 - 1 |

To determine if *MGD1* and *MGD2* also contributed to MG metabolism, we tested whether the *mgd1∆*, *mgd2∆*, or *mgd1∆*/*mgd2∆* mutants were deficient in utilizing MG as a sole carbon source. In minimal YNB medium with 5 mM glucose, none of the mutants displayed a significant difference in final OD_600_ relative to the WT (**Fig. S2A**). With 5 mM MG as the sole carbon source, neither single mutant exhibited a significant defect in growth, but the *mgd1∆*/*mgd2∆* displayed a 26.8% reduction in yield (p < 0.05) relative to the WT (**Fig. S2B**). Overall, the results in Fig. 2 and **Fig. S2** suggest that both *MGD1* and *MGD2* play a role in the detoxification and metabolism of MG.

### Mrr1 regulates expression of *MGD1*, and *MGD2* is not highly expressed under standard conditions

In Demers et al (7), we reported an RNA-seq analysis that showed that clinical isolates with constitutive Mrr1 activity had higher levels of *MGD1* and *MGD2* expression than strains with low basal Mrr1 activity. Furthermore, analyses of *C*. *albicans*, *C*. *parapsilosis*, and an independent collection of clinical *C. lusitaniae* isolates also found that *GRP2/MGD1* gene expression was elevated in azole-resistant strains with gain of function mutations in Mrr1 (13–18). Mrr1 was required for resistance to MG, as the S18 *mrr1∆* mutant was significantly more sensitive to 15 mM MG than the WT (Fig. 3A). Similar results were obtained in strain L17 thus validating the mutant phenotype (Fig. 3B). In either background, there was no difference in growth between the WT and *mrr1Δ* mutant in the absence of MG (Fig. 3A-B). Additionally, we found that the *mrr1∆* mutant had a 31.8% lower yield relative to the parental strain in MG as a sole carbon source with no defects in growth on glucose (**Fig. S2**). Thus, knocking out *MRR1* had similar effects on growth with MG as knocking out *MGD1* and *MGD2*.

**Fig. 3.**
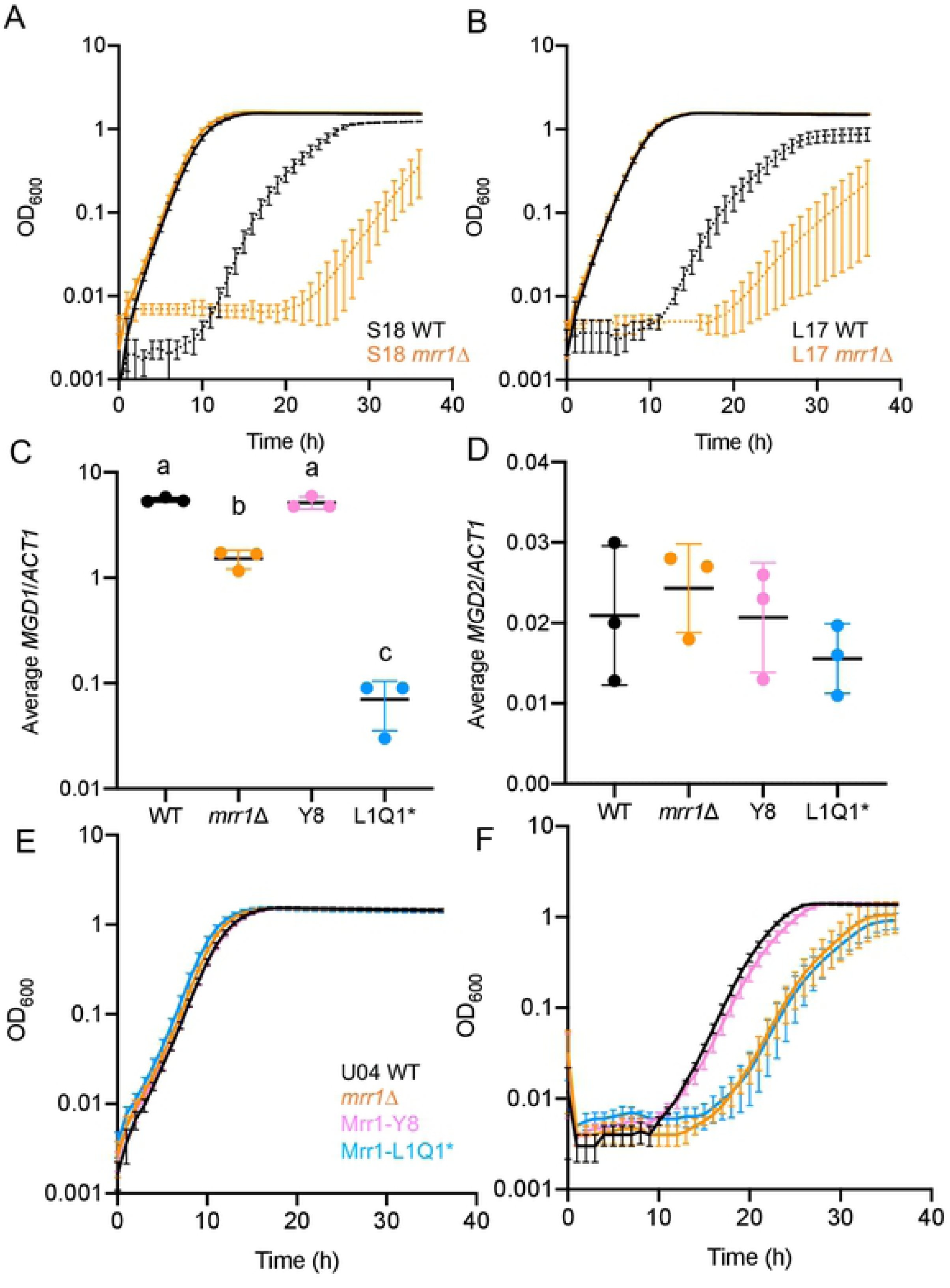
Mrr1 regulates MG resistance and basal expression of *MGD1* but not *MGD2*. **(A-B)** Growth curves of *C. lusitaniae* S18 **(A)** and L17 **(B)** wild type (black) and *mrr1∆* (orange) in YPD alone (solid) or with 15 mM MG (dashed). One representative experiment out of three independent experiments is shown. **(C-D)** Expression of *MGD1* **(C)** and *MGD2* **(D)** in *C. lusitaniae* U04 (black), U04 *mrr1∆* (orange), U04 *mrr1∆* + *MRR1-Y8* (pink) and U04 *mrr1∆* + *MRR1-L1Q1\** (light blue). Three-way ANOVA was used for statistical evaluation. **(E-F)** Growth curves of *C. lusitaniae* U04 (black), U04 *mrr1∆* (orange), U04 *mrr1∆* + *MRR1-Y8* (pink) and U04 *mrr1∆* + *MRR1-L1Q1\** (light blue) in YPD alone **(E)** or with 15 mM MG **(F)**. One representative experiment out of three independent experiments is shown.

To more directly assess whether Mrr1 controls expression of *MGD1* and *MGD2*, we analyzed expression of *MGD1* and *MGD2* in an existing set of isogenic strains that differed only by which *MRR1* allele was present at the native locus. The naturally-occurring *MRR1* alleles included in this set included Mrr1-Y813C (Y8), the constitutively-active native allele of U04, and Mrr1-L1191H + Q1197* (L1Q1*), which is a prematurely truncated allele that has low activity and confers sensitivity to FLZ (Table 1) (7). Each allele was introduced into an *mrr1∆* derivative of strain U04. The *mrr1Δ* mutant had significantly lower levels of basal expression of *MGD1* relative to the U04 WT, and complementation with the native Mrr1-Y8 restored expression of *MGD1* to WT levels (Fig. 3C). Consistent with Mrr1 acting as a positive regulator of *MGD1*, the strain with low activity Mrr1-L1Q1* had low *MGD1* expression (Fig. 3C). The difference between the *mrr1∆* and the Mrr1-L1Q1* strain suggests complex roles for low activity Mrr1, although more work is needed to elucidate the mechanism by which this occurs. *MGD2* levels were lower than *MGD1* by 10 - 100-fold, as judged by comparison to a reference transcript and a standard curve determined for each primer set (see Methods). Surprisingly, *MGD2* levels were not different across the U04 strains with different Mrr1 variants under these experimental conditions (Fig. 3D). In line with the transcriptional differences between strains with high Mrr1 activity (U04 WT and U04+Mrr1-Y8) and strains with low or no Mrr1 activity (*mrr1∆* or Mrr1-L1Q1*), we found that strains with low Mrr1 activity grew less well with MG with no differences in control conditions (Fig. 3E-F). Together, these data strongly indicate that *MGD1* is regulated by Mrr1; the conditional role of Mrr1 in the control of *MGD2* expression is addressed further below.

### Exogenous MG induces Mrr1-regulated genes and increases FLZ resistance

Given that MG induces levels of MG reductase in *S. cerevisiae* (40) along with our observations that *MGD1* and *MGD2* are involved in detoxification and metabolism of MG (Fig. 2 and **S2B**), we hypothesized that MG may induce expression of these two genes in *C. lusitaniae*. As shown in Fig. 4A-B, 5 mM MG significantly induced expression of *MGD1 and MGD2* by 2- and 16-fold, respectively at 15 minutes and 30 minutes. Expression of *MGD1* remained around 2-fold higher than the control by 30 minutes. Although expression of both genes appeared elevated after 60 minutes of MG exposure, the difference relative to the basal expression was only significant for *MGD1*, indicating that expression of these genes returns to basal levels over time. We also tested whether MG would also induce expression of *MDR1*, another gene regulated by Mrr1, and found that 5 mM MG induced *MDR1* expression with a 6-fold within 15 and 30 minutes (Figure 4C). As with *MGD1* and *MGD2*, relative *MDR1* levels decreased by 60 minutes. These results suggest that MG induces a rapid transcriptional response in *C. lusitaniae*, which includes upregulation of multiple Mrr1-regulated genes.

**Fig. 4.**
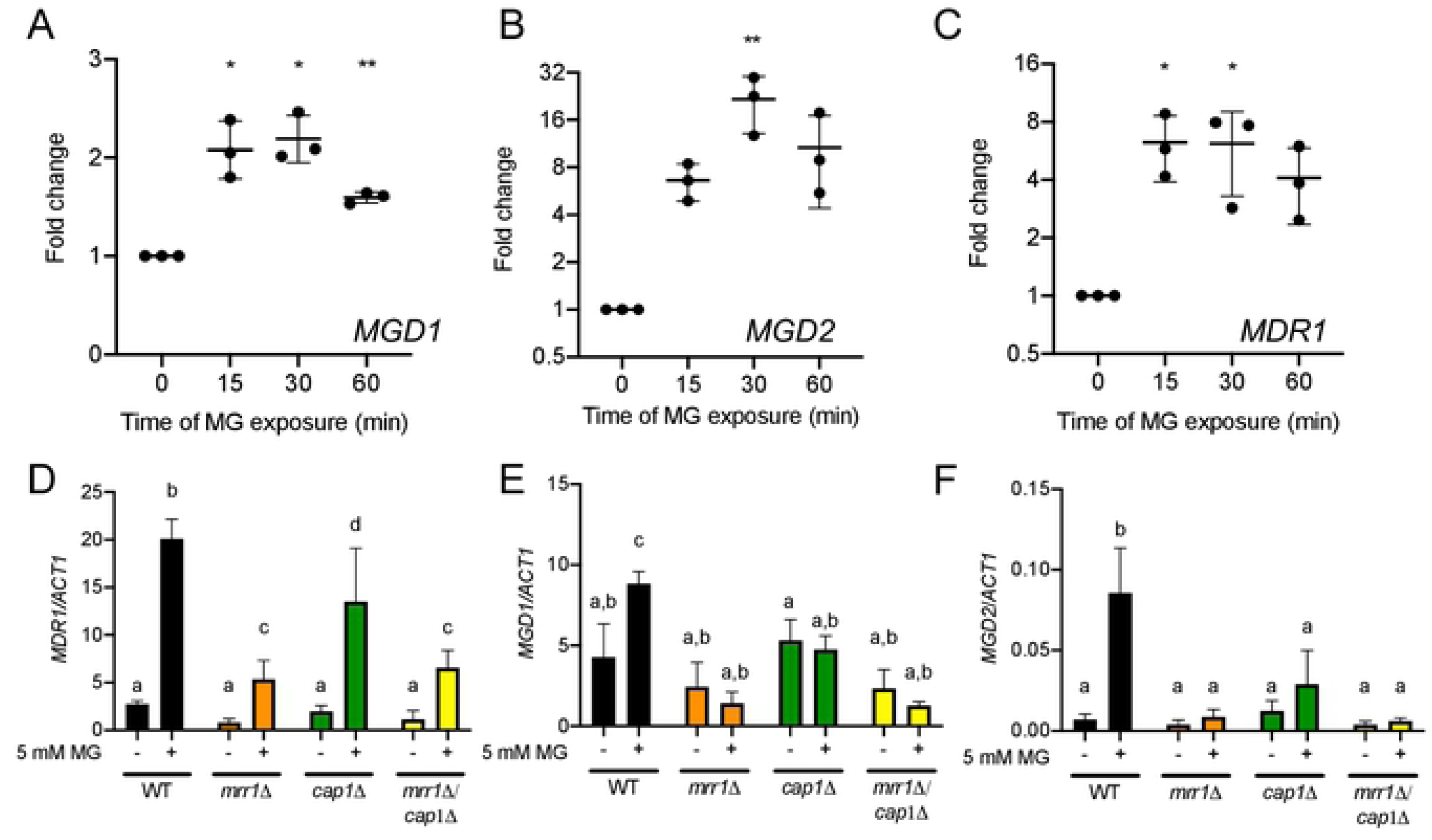
Expression of *MGD1*, *MGD2*, and *MDR1* in response to MG. **(A-C)** *C. lusitaniae* S18 was grown to exponential phase and treated with 5 mM MG for the time indicated prior to analysis of *MGD1* **(A)**, *MGD2* **(B)**, and *MDR1* **(C)** transcript levels by qRT-PCR. Transcript levels are normalized to levels of *ACT1* and presented as ratio at each time point relative to 0 min for three independent experiments. One-way ANOVA was used for statistical evaluation: * p < 0.05, ** p < 0.01. **(D-F)** *C. lusitaniae* S18 wild type (black) and *mrr1∆* (orange), *cap1∆* (green), and *mrr1∆*/*cap1∆* (yellow) mutants were grown to exponential phase and treated with 5 mM MG for 15 minutes prior to analysis of *MDR1* **(D)**, and *MGD1* **(E)**, and *MGD2* **(F)** transcript levels by qRT-PCR. Transcript levels are normalized to *ACT1*. Data shown represent the mean ± SD for three independent experiments. Three-way ANOVA was used for statistical evaluation.

### Mrr1 and Cap1 play a role in the transcriptional response to MG

Mrr1 induces *MDR1* in response to benomyl and hydrogen peroxide (H_2_O_2_) in *C. albicans* (13) through an unknown mechanism. We hypothesized that Mrr1 may also mediate the induction of *MDR1*, *MGD1*, and *MGD2* in response to MG in *C. lusitaniae*. We also analyzed the role of the bZIP transcription factor Cap1 in the MG response as in *C. albicans*, Cap1 is also required for upregulation of *MDR1* in response to H_2_O_2_ (13), and ChIP analyses found Cap1 bound to the promoters of *MDR1* and *GRP2*/*MGD1* (41). Furthermore, in *S. cerevisiae*, MG directly modifies the Cap1 ortholog Yap1 by reversibly oxidizing cysteines, facilitating nuclear localization (34).

To determine whether *C. lusitaniae MRR1* and/or *CAP1* (*CLUG_02670*) were required for the induction of transcription of *MDR1* in response to MG, we repeated our transcriptional induction experiment with the *mrr1*∆, *cap1*∆, and *mrr1*∆/*cap1*∆ mutants in addition to S18 WT. In the absence of MG, there were no significant differences in *MDR1* expression between strains (Fig. 4D) though *MDR1* expression trended lower in the *mrr1*∆ and *mrr1*∆/*cap1*∆ mutants. The WT showed a strong (~10-fold), significant increase in *MDR1* expression after 15 minutes. Much lower levels of induction were observed in the *mrr1∆, cap1∆*, and *mrr1∆*/*cap1∆* mutants (Fig. 4D). As in S18, MG-treated L17 *mrr1*∆ and *cap1∆* had a significantly lower *MDR1*:*ACT1* ratio than the MG treated WT, supporting the model that Mrr1, and to a lesser extent Cap1, participated in the transcriptional induction of *MDR1* by MG (**Fig. S3A**). Using the same sample sets, we found that S18 WT exhibited an approximately two-fold and 12-fold increases in *MGD1* and *MGD2* expression, respectively, in response to MG, while none of the mutants had a significant change in *MGD1* or *MGD2* expression (Fig. 4E-F) suggesting that, in response to MG, MG reductases are co-regulated with *MDR1* through Mrr1 and Cap1. Like the mutants lacking Mrr1, the *cap1∆* mutant was also defective in growth in YPD + 15 mM MG (**Fig. S4**) providing further evidence for Mrr1-Cap1 co-regulation of MG-detoxification.

### MG stimulates growth in FLZ in an Mrr1- and Mdr1-dependent manner

Due to the induction of *MDR1* expression by MG, we hypothesized that MG could increase *MDR1*-dependent FLZ resistance in *C. lusitaniae*. We performed growth kinetics with MG and FLZ on S18 WT and its *mrr1∆*, *cap1∆*, *mrr1∆*/*cap1∆*, and *mdr1*∆ derivatives. For each strain, we used FLZ at a concentration that allowed for minimal growth, which for most strains was half of the MIC (Table 1). While 5 mM MG did not affect the growth of S18 WT (Fig. 5) or any of the mutants (Fig. 5B) in the absence of FLZ, MG drastically improved growth in the presence of FLZ compared to FLZ alone (Fig. 5). The increased FLZ resistance in the presence of MG was dependent on *MDR1*. The loss of *MRR1* strongly and significantly reduced the benefits of MG on growth with FLZ, and the effects of MG were completely absent in the *mrr1*∆/*cap1*∆ double mutant (Fig. 5B). We repeated these growth assays in the L17 background with knockout mutants of *MRR1*, *CAP1*, and *MDR1*. Strain L17 was not amenable to the construction of double mutants due to unknown factors, and thus the *mrr1∆cap1∆* double was not analyzed. Similar to S18, the parental isolate exhibited robust stimulation of growth in the presence of MG (**Fig. S3B**) and the *mdr1*∆ mutant lost most of the stimulation seen in the unaltered parent (**Fig. S3C**). However, the effects of knocking out *MRR1* or *CAP1* differed in L17 in terms of the relative contributions of each transcription factor (**Fig. S3C**). Overall, Mrr1 and Cap1 may be differentially important for the MG response based on genetic backgrounds or other assay variables, but that they both contribute to FLZ resistance in the presence of MG.

**Fig. 5.**
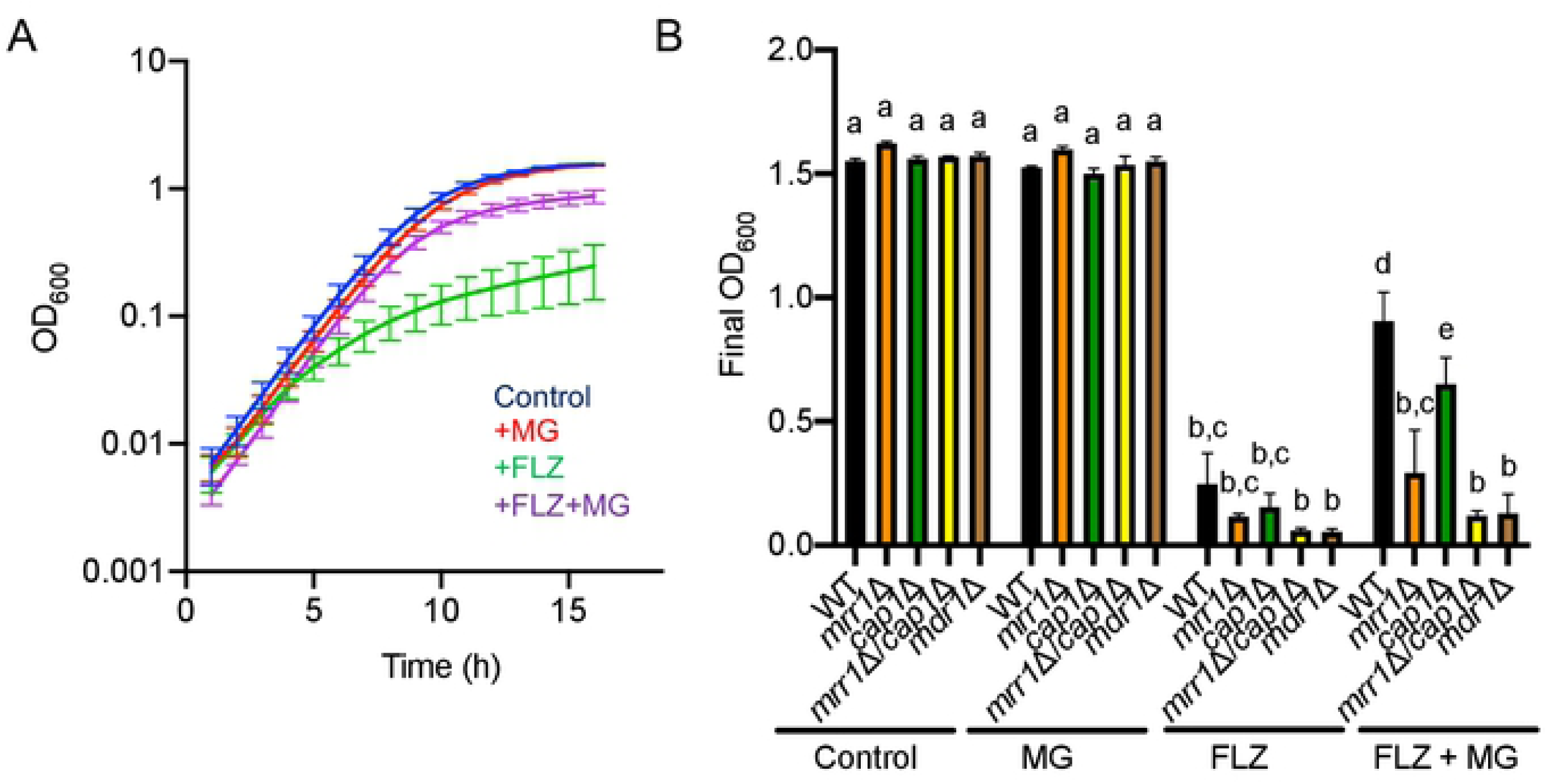
MG increases fluconazole (FLZ) tolerance that is dependent on *MRR1* and *MDR1* in isolate S18. **(A)** *C. lusitaniae* S18 grown in YPD alone (blue), or with 5 mM MG (red), FLZ (1/2 MIC) (green), or FLZ + 5 mM MG (light purple). **(B)** Final OD_600_ at 16 hours of growth for each indicated strain in YPD alone or with 5 mM MG and/or FLZ. Data shown represent the mean ± SD from three independent experiments. Three-way ANOVA was used for statistical evaluation.

### Constitutively active Mrr1 results in greater stimulation of growth in FLZ than truncated Mrr1

Given our discovery of repeated selection for Mrr1 variants with constitutive activity within a chronic *C. lusitaniae* lung infection population, we sought to determine if higher basal Mrr1 activity enhanced the stimulation of FLZ resistance by MG. We compared the effects of MG on growth in the presence of FLZ for *C. lusitaniae* strains S18 and L17, which both express a constitutively active Mrr1 variant (Mrr1-H467L; H4) to previously published strains U05 and L14, which express the low activity Mrr1-L1Q1* variant. Strains S18 and L17 showed a greater improvement in growth with FLZ upon inclusion of MG than strains U05 and L14 with low activity Mrr1 (Fig. 6A). In a direct comparison of final yield, there were no differences in growth among strains in YPD alone, 5 mM MG, or in FLZ at half of each strain’s MIC (Fig. 6B). FLZ significantly inhibited growth of all strains relative to the YPD controls. In medium with FLZ and MG, all four strains have a significantly higher OD_600_ than in FLZ alone, but S18 and L17 grew to a significantly higher OD_600_ than U05 and L14 (Fig. 6B). These results suggest that constitutively active Mrr1 variants can enhance stimulation of Mrr1 by MG.

**Fig. 6.**
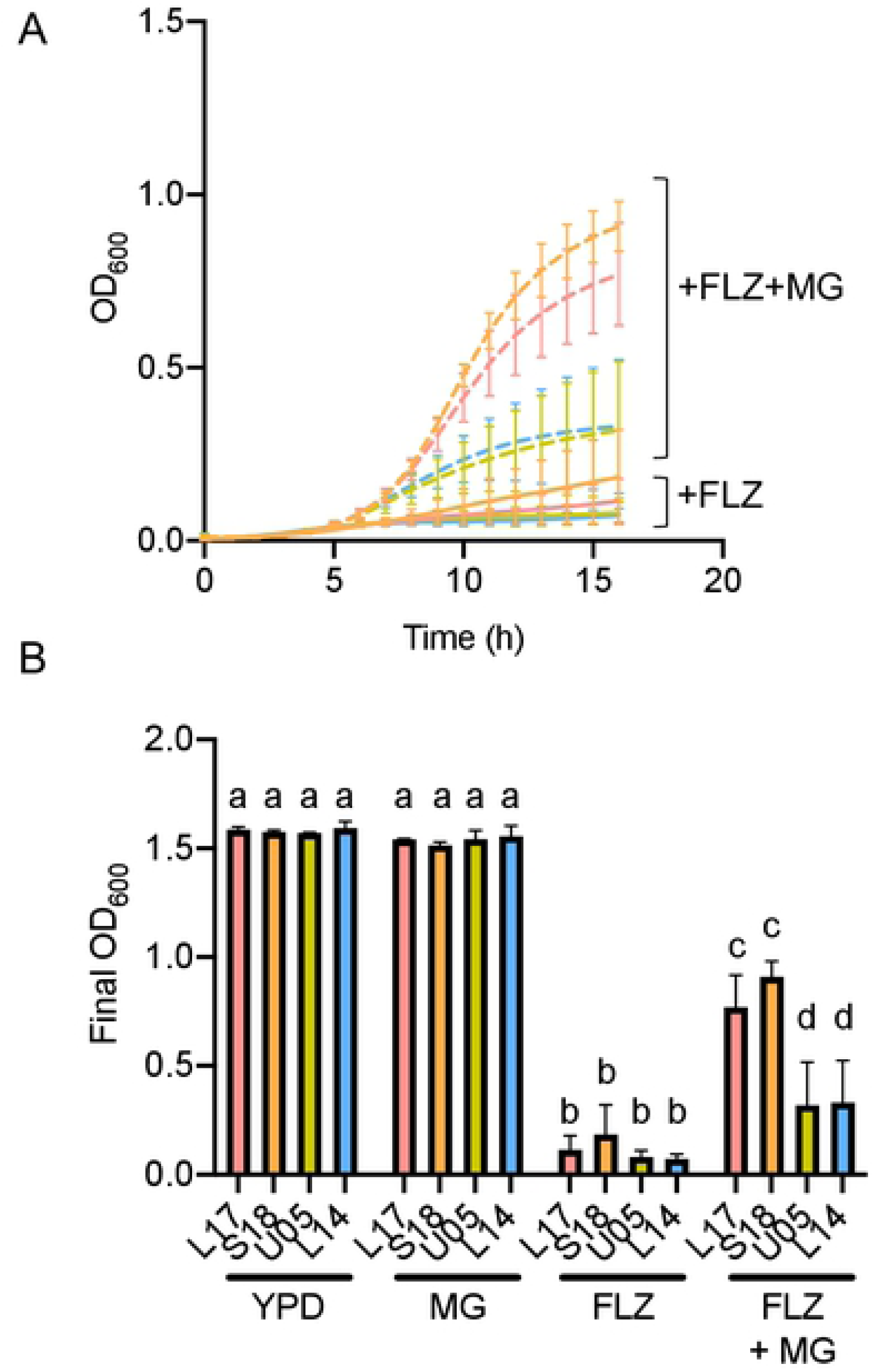
Constitutively active Mrr1 leads to a greater increase in stimulation of growth in FLZ by MG compared to truncated Mrr1. *C. lusitaniae* isolates U05 (green), L14 (blue), L17 (pink), and S18 (orange) were grown in YPD with 5 mM MG and/or FLZ. **(A)** Growth kinetics of isolates U05, L14, S18, and L17 in FLZ with (dashed lines) or without (solid lines) 5 mM MG. **(B)** Final OD_600_ at 16 hours of growth in each condition. Data shown represent the mean ± SD from three independent experiments. Three-way ANOVA was used for statistical evaluation.

### Absence of *GLO1* causes increased sensitivity to MG and increased resistance to FLZ

The experiments above focused on the effects of exogenous MG. Because MG is a highly reactive molecule and thus actual exposure concentrations are difficult to determine, we sought to assess the effects of endogenous MG on Mrr1 activation and FLZ resistance. The glyoxalase pathway, which consists of the glutathione-dependent enzymes Glo1 and Glo2, is widely recognized as a major mechanism for MG detoxification in eukaryotic cells (see Fig. 1) (42), and we speculated that knocking out *GLO1* would increase both MG sensitivity, due to decreased MG detoxification capacity, and increased levels of endogenous MG. In strain S18, we knocked out *GLO1* (*CLUG_04105*) and found that the *glo1*∆ mutant was completely inhibited by 15 mM MG (Fig. 7A), whereas the *mgd1*∆, *mgd2*∆, and *mgd1*∆/*mgd2*∆ mutants were still able to grow at this concentration (Fig. 2B-D). Because all cells naturally produce MG as a metabolic byproduct, and cells lacking *GLO1* accumulate more intracellular MG (34, 43), we were interested in whether the S18 *glo1*∆ mutant was more resistant to FLZ than its parent. Although S18 *glo1*∆ had the same FLZ MIC as wild type S18 (Table 1) and similar growth kinetics in YPD (Fig. 7A), the *glo1∆* strain grew substantially better in FLZ compared to the S18 WT (Fig 7B). These data suggest that endogenous MG, at concentrations that do not inhibit growth in YPD, confer some resistance to FLZ. More work is needed to better understand the role of endogenous MG in *Candida* species and whether changes in intracellular glutathione would influence intracellular MG levels.

**Fig 7.**
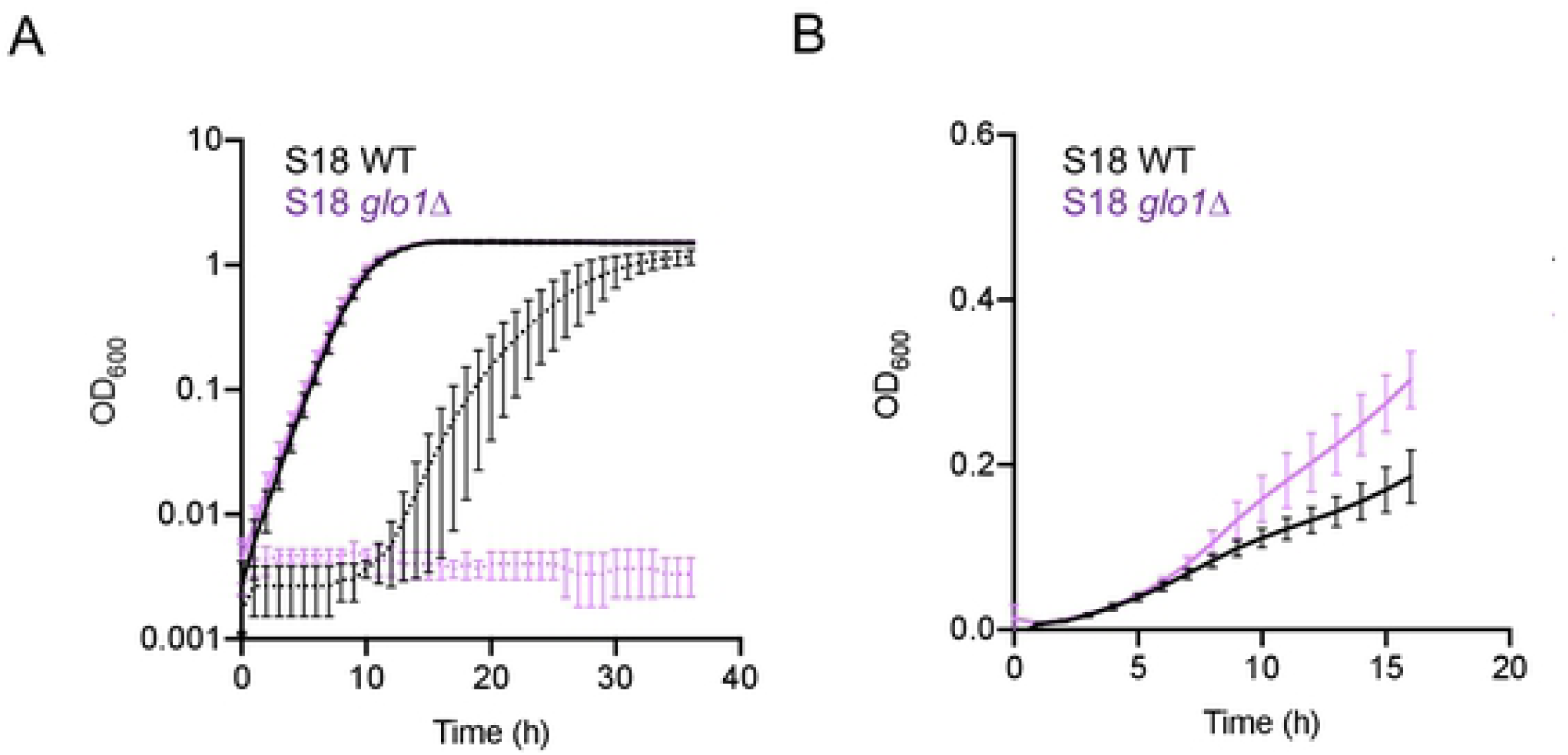
Knocking out *GLO1* leads to increased sensitivity to MG and increased resistance to FLZ. *C. lusitaniae* S18 wild type (black) and *glo1∆* (light purple) were grown in YPD with (dashed) or without (solid) 15 mM MG **(A)** or YPD with 8 µg/mL FLZ **(B)**. Data shown represent the mean ± SD from three independent experiments.

### *C. lusitaniae* is more resistant to MG than many other *Candida* species, but some strains of other species can exhibit induction of FLZ resistance by MG

We sought to determine if MG induced azole resistance in other *Candida* species. To do so, we first we assessed MG sensitivity across strains and species. As controls for the effects of MG, we included *C. lusitaniae* isolate S18 and its *glo1*∆ mutant. Of the other *C. lusitaniae* isolates that we tested, all grew at least as well as S18 on MG with one strain, Y533, growing slightly better (Fig. 8A). We observed notable heterogeneity in MG sensitivity among the tested *Candida auris* strains: CAU-01 was completely inhibited by 15 mM MG, while *C. auris* CAU-05 grew about as well as *C. lusitaniae* strain S18 (Fig. 8A). The other three *C. auris* strains exhibited intermediate phenotypes. *Candida albicans* strain SC5314 displayed weak growth on 15 mM MG, but the two clinical *C. albicans* isolates F2 and F5 appeared to be more resistant (Fig. 8A). The single *Candida guillermondii* strain and two *Candida glabrata* strains we tested were all highly sensitive to MG (Fig. 8A). Likewise, all three *Candida parapsilosis* strains failed to grow on 15 mM MG (Fig. 8A). Finally, the three *Candida dubliniensis* strains grew moderately well on MG, with no visible differences between strains (Fig. 8A). Overall, the results in Fig. 8A, using a limited number of strains, suggest that MG sensitivity varies across *Candida* species and strains.

**Fig 8.**
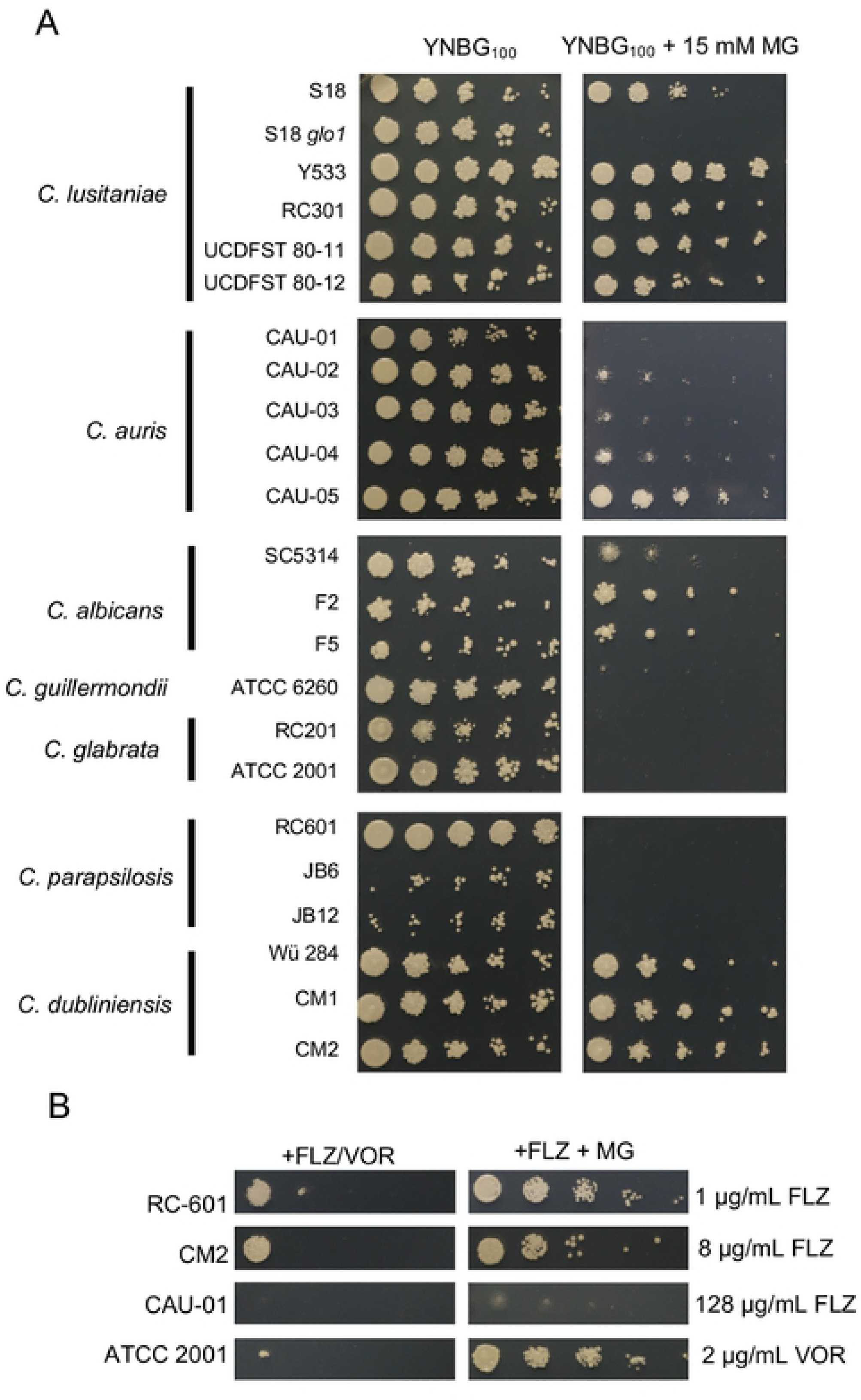
Effect of MG on other *Candida* species and strains. **(A)** Serial 1:10 dilutions of each *Candida* strain were spotted onto YNBG100 without or with 15 mM MG, then grown at 37°C for two days. One representative out of three independent experiments is shown. **(B)** Growth on FLZ (at ½ MIC concentration as indicated) in the absence and presence of MG across different *Candida* species and strains. One representative experiment out of two independent experiments is shown. UCDFST, Phaff Yeast Culture Collection, Food Science and Technology, University of California Davis; ATCC, American Type Culture Collection.

To investigate whether MG induces azole resistance in other *Candida* species, we performed a serial dilution using varying concentrations of FLZ with or without 3 mM MG. We used a lower concentration of MG due to the higher MG sensitivity of some species compared to *C. lusitaniae*. As shown in Fig. 8B, *C. parapsilosis* RC-601 and *C. dubliniensis* CM2 displayed a striking increase of growth on FLZ with MG. *C. auris* CAU-01 demonstrated a more subtle increase in growth (Fig. 8A). Finally, *C. glabrata* ATCC 2001 exhibited a striking increase of growth on voriconazole (VOR) with MG (Fig. 8A); we used VOR because this strain was not appreciably inhibited by FLZ at any of the concentrations tested. All other strains not shown in Fig. 8B did not demonstrate visible stimulation of growth on FLZ by MG under the tested conditions. These results suggest that MG stimulation of azole resistance is not exclusive to *C. lusitaniae*, but not every strain within a species can be stimulated under the conditions tested.

## Discussion

Endogenous inducers of the important azole resistance associated transcriptional regulator Mrr1 activity are not yet well understood. In *C. albicans*, *MDR1* expression can be induced by a variety of toxic compounds, such as methotrexate, 4-nitroquinoline-N-oxide, *o*-phenanthroline, benomyl, diethyl maleate, diamide and H_2_O_2_, (12, 13, 44), many of which induce oxidative stress. However, of these, only H_2_O_2_ is known to be present *in vivo* and to our knowledge endogenous H_2_O_2_ production has not been shown to cause increased Mrr1 activity or FLZ resistance. In this work, through the fact that Mrr1 regulates the expression of *MGD1* (Fig 3C) and *MGD2* (Fig. 4D-E), which encode enzymes necessary for MG resistance (Fig. 2) and catabolism (**Fig. S2**), we found that exogenous (Fig. 5 and **S3**) and endogenous (Fig. 7B) MG enhanced resistance to FLZ in clinical isolates of the opportunistic pathogen *C. lusitaniae* and other *Candida* species (Fig. 8B). The increased resistance to FLZ is largely dependent on induction of *MDR1* (Fig. 5B and **S3C**) but may involve other MG-induced factors such as the co-regulated *FLU1*, a gene which encodes another efflux pump that has been associated with resistance against FLZ, cycloheximide, and mycophenolic acid in *C. albicans* (45, 46). Strains with an *MRR1* allele with a gain-of-function mutation had a larger MG-dependent increase in growth in FLZ than strains with low activity Mrr1 (Fig. 6), leading us to suggest that strains with elevated MICs might have even greater responses to host MG. MG is an attractive inducer of Mrr1 activity because it is higher in many patients who are at greater risk for colonization or infection by *Candida* species relative to the general population.

Importantly, diabetes, which is associated with elevated serum MG (21–23), is also considered a risk factor for colonization and infection by a variety of *Candida* species (reviewed in (47)). Kidney failure, another disease in which MG levels are higher than healthy controls (25–28), is also associated with increased risk of *Candida* infection (48, 49). Thus, our demonstration of the induction of azole resistance by MG could be an important step toward understanding and preventing treatment failure in populations who are susceptible to *Candida* colonization and infection.

Although the serum concentrations of MG reported in humans (50, 51) are lower than those used here, MG is highly reactive and thus it is challenging if not impossible to assess local MG concentrations near sites of its production. Neutrophils may produce MG in response to pathogens; a study in Group A *Streptococci* found that deletion of the glyoxalase I gene *gloA* rendered cells more susceptible to neutrophil killing and this survival defect was rescued with addition of a myeloperoxidase inhibitor (29). Furthermore, MG is substantially elevated in diabetes (21–23, 52), uremia (25–28), or sepsis (24), and it is not currently known how sustained exposure to MG affects microbes that reside in patients with these or similar diseases. Finally, metabolism of different carbon sources *in vivo* may result in overproduction of MG by host cells and/or microbial cells. For instance, fructose is elevated in individuals with diabetes and metabolic syndrome, and evidence suggests that fructose leads to greater accumulation of MG in certain mammalian tissues (53, 54).

The mechanisms by which MG activates transcription will be the subject of future work. Multiple studies have established that Mrr1 and Cap1 cooperate to regulate expression of certain genes in *C. albicans*, including *MDR1* (12, 13). Other studies have demonstrated a role for Cap1 itself in regulating expression of *MDR1* (41, 55). This information, along with the published observation that MG activates Yap1 (34), the *S. cerevisiae* homolog of Cap1, led us to hypothesize that Mrr1 and Cap1 cooperate to upregulate expression of *MDR1*, *MGD1*, and *MGD2* in the presence of MG. Our studies suggest that *CAP1* may be involved in but is not required for the response of *C. lusitaniae* to MG which results in elevated resistance to FLZ. Unlike *MGD1* and *MGD2* which showed no induction in the *mrr1∆*/*cap1∆* mutant (Fig. 4E-F), *MDR1* still showed some residual induction in response to MG (Fig 4D). One possible explanation for this unexpected result is that Upc2, which has been shown to be a minor regulator of *MDR1* expression in *C. albicans* (13, 56), may be regulating expression and/or induction of *MDR1* in the absence of *MRR1* and *CAP1.* In *C. albicans*, Upc2 requires functional Mrr1 to upregulate *MDR1* expression (13), but this may not be the case in *C. lusitaniae*. Another transcription factor involved in *MDR1* expression in *C. albicans* is Mcm1, which is required for induction of *MDR1* by benomyl and by hyperactive Mrr1, but not induction by H_2_O_2_ (12). Finally, the Swi/Snf chromatin remodeling complex is also required for *MDR1* overexpression mediated by hyperactive Mrr1 in *C. albicans* (11). Therefore, candidates for direct activation by MG in *C. lusitaniae* include Cap1, the homolog of which, in *S. cerevisiae*, is reversibly modified at cysteine residues, disrupting its interaction with a nuclear exportin (34); Mrr1, which also contains many cysteine residues near the C-terminal portion; as well as Upc2, Mcm1, and the Swi/Snf complex.

*GLO1* encodes a glutathione-dependent glyoxalase that also catabolizes MG and is a major mechanism for detoxification of MG in eukaryotes; loss of *GLO1* leads to elevated intracellular MG in *S. cerevisiae* (34, 43). Here, we show that a *C. lusitaniae glo1* mutant is highly sensitive to exogenous MG (Fig. 7A), and in the absence of exogenous MG it demonstrates improved growth in FLZ when compared to its unaltered parental isolate S18 (Fig. 7B). This suggests that the buildup of intracellular MG resulting from *GLO1* deficiency may be sufficient to induce FLZ resistance. The glyoxalase system requires reduced glutathione (GSH) to function, so it is possible that oxidants encountered *in vivo* may deplete GSH and lead to increased intracellular MG. In fact, GSH levels are reduced by chronic stress or in chronic infections accompanied by oxidative stress, such as cystic fibrosis (57, 58). It is also worth noting that diethyl maleate, a compound that induces *MDR1* expression in *C. albicans* (44), is commonly used in laboratory studies to deplete GSH (59–63).

We were struck by the differences between strains and species in both MG resistance and MG-stimulation of FLZ resistance. *C. lusitaniae* isolates were overall quite resistant to MG compared to many other *Candida* species (Fig 8A). Interestingly, UCDFST 80-11 and UCDFST 80-12, which are closely related and were both isolated from rotting cactus, grow about as robustly on 15 mM MG as the three clinical isolates (S18, Y533, and RC-301), indicating that, among these samples, there is not a clear phenotypic difference in MG sensitivity between environmental isolates of *C. lusitaniae* versus those obtained from a human host. Under ambient conditions, MG reaches a concentration of 30-75 µM in various plant species and can increase 2- to 6-fold during stressful conditions such as drought or high salinity (64). Thus, it is possible that *C. lusitaniae*, which is found predominantly in the environment rather than in the human microbiota, encounters substantial amounts of MG while living on plant hosts, leading to a general, species-wide MG resistance. The highly drug-resistant *C. auris* is closely related to *C. lusitaniae* (65) and likewise has two putative MG reductase genes with high identity (Fig. 2A). Therefore, we expected *C. auris* to be approximately as resistant to MG as *C. lusitaniae*. However, we observed marked heterogeneity among the five *C. auris* strains tested (Fig. 8A), with CAU-05 demonstrating growth on MG similar to the *C. lusitaniae* strains. CAU-01 was highly sensitive to MG and it is interesting to note that this strain is more sensitive to FLZ, H_2_O_2_, and neutrophil killing compared to other *C. auris* strains (66). A better understanding of how the *in vivo* environment may influence drug resistance of clinical isolates may enable a more strategic evaluation of which drugs or drug combinations are most likely to be successful.

## Methods

### Generation of MG reductase phylogenetic tree

Orthologs of Gre2 across *Candida* species were identified in FungiDB (https://fungidb.org) and selected for a protein Clustal Omega multiple sequence alignment. The resulting alignment was then used to generate a phylogenetic tree using the Interactive Tree of Life (ITOL) tool (https://itol.embl.de).

### Strains, media, and growth conditions

The sources of all strains used in this study are listed in **Table S1**. All strains were stored long term in a final concentration of 25% glycerol at −80°C and freshly streaked onto YPD agar (10 g/L yeast extract, 20 g/L peptone, 2% glucose, 1.5% agar) once every seven days. Unless otherwise noted, all overnight cultures were grown in 5 mL YPD liquid medium (10 g/L yeast extract, 20 g/L peptone, 2% glucose) on a rotary wheel at 30°C.

### Mutant construction

Mutants were generated using an expression-free CRISPR-Cas9 method, as previously described (67). In brief, cultures were grown to exponential phase in 50 mL YPD, then washed and incubated in TE buffer and 0.1 M lithium acetate at 30°C for one hour. Dithiothreitol was added to a final concentration of 100 mM and cultures were incubated for an additional 30 minutes at 30°C. Cells were washed and resuspended in 1 M sorbitol before being transferred to electroporation cuvettes. To each cuvette was added 1.5 µg of knockout construct and Cas9 ribonucleoprotein containing crRNA specific to the target gene. Following electroporation, cells were allowed to recover in YPD at 30°C for four to six hours. Cells were then plated on YPD agar with selection and incubated at 30°C for two days. Primers used to create knockout constructs are listed in **Table S2**.

### Minimum Inhibitory Concentration (MIC) Assay

MIC assays for FLZ were performed as described in (7). In brief, overnight cultures were diluted to an OD_600_ of 0.1 in 200 µL dH_2_O and 60 µL of each dilution were added to 5 mL Roswell Park Memorial Institute (RPMI) Medium, pH 7.0. FLZ was serially diluted across a clear, flat-bottom 96-well plate (Falcon) from 128 µg/mL down to 0.25 µg/mL in RPMI, pH 7.0. To each well was added 100 µL of cell suspension in RPMI. Upon addition of cells, the final concentration of FLZ ranged from 64 µg/mL to 0.125 µg/mL. Plates were incubated at 35°C and scored for growth at 24 and 48 hours; the results reported in Table 1 are at 48 hours of growth.

### Growth Kinetics

*C. lusitaniae* cultures were grown overnight, diluted 1:50 into 5 mL fresh YPD, and grown for four to six hours at 30°C. After washing, the cultures were diluted to OD_600_ 1 in 200 µL dH_2_O. Each inoculum was prepared by pipetting 60 µL of OD_600_ 1 dilution into 5 mL YPD. Clear 96-well flat-bottom plates (Falcon) were prepared by adding 100 µl per well YPD or YPD with MG and/or FLZ at twice the desired final concentrations. 100 µL of inoculum was added to each row of the plate. Each plate was set up in technical triplicate for each strain and condition. The plates were incubated in a Synergy Neo2 Microplate Reader (BioTek) to generate a kinetic curve. The plate reader protocol was as follows: heat to 37°C, start kinetic, read OD_600_ every 60 minutes for 16 or 36 hours, end kinetic.

### Spot Assays

*Candida* cultures were grown overnight, diluted 1:50 into 5 mL fresh YPD, and grown to exponential phase at 30°C. Cultures were diluted to OD_600_ 1 in 200 µL dH_2_O. Each strain was then serially diluted 1:10 until a final OD_600_ of approximately 1 x 10^−6^. 5 µL of each dilution was spotted onto the specified medium. Plates were incubated at 37°C for two days, and then photographed.

### Quantitative Real-Time PCR

*C. lusitaniae* cultures were grown overnight, diluted 1:50 into 5 mL fresh YPD, and grown for four hours at 30°C. Control cultures were harvested at this point and MG was added to a final concentration of 5 mM to all other cultures, which were returned to 30°C. Cultures were then harvested after 15, 30, or 60 minutes. To harvest, 2 mL of culture were spun in a tabletop centrifuge at 13.2 x *g* for 5 min and supernatant was discarded. RNA isolation, gDNA removal, cDNA synthesis, and quantitative real-time PCR were performed as previously described (7). Primers are listed in **Table S2**.

### Statistical Analysis

All graphs were prepared with GraphPad Prism 8.3.0. A linear model was used to estimate the effect of strain differences and exposure to MG, including the interaction between strain and exposure to MG. Linear models were run in R. One- and three-way analysis of variance (ANOVA) tests were performed in Prism. All p values are two-tailed and p < 0.05 was interpreted to be statistically significant.

## Acknowledgements

We thank Thomas Hampton for assistance with the linear model used to analyze the data in Fig. 2. We thank Theodore White for providing *C. lusitaniae* strain Y533; Richard Calderone for providing *C. guillermondii* strain ATCC 6260, *C. glabrata* strain RC-201, *C. lusitaniae* strain RC-301, and *C. parapsilosis* strain RC-601; Lawrence Myers for providing *C. dubliniensis* strain Wü284; the FDA-CDC Antimicrobial Resistance Isolate Bank for providing *C. auris* strains CAU-01, CAU-02, CAU-03, CAU-04, and CAU-05; the American Type Culture Collection (ATCC) for providing *C. glabrata* strain ATCC 2001; Kyria Boundy-Mills for *C*. *lusitaniae* strains UCDFST 80-11 and UCDFST 80-12; Joachim Morschhäuser for *C. albicans* strains F2 and F5 as well as *C. dubliniensis* strains CM1 and CM2, and Isabel Miranda for *C. parapsilosis* strains JB6 and JB12. We acknowledge the Shared Resources facility bioMT Molecular Interactions & Imaging Core (MIIC) at the Norris Cotton Cancer Center at Dartmouth.

## Footnotes

### Author contributions

ARB, EGD, and DAH conceived and designed the experiments and wrote the paper. ARB and EGD performed the experiments. ARB, EGD, and DAH analyzed the data.

### Funding

This study was supported by grants R01 5R01AI127548 to DAH, AI133956 to EGD, STANTO19R0 to CFF RDP, P30-DK117469 to DartCF, and P20-GM113132 to BioMT. The Cystic Fibrosis Research Development Program and NIGMS of the NIH as P20GM103413 supported the Translational Research Core at Dartmouth. Sequencing services and specialized equipment were provided by the Molecular Biology Shared Resource Core at Dartmouth, NCI Cancer Center Support Grant 5P30 CA023108-41. The content is solely the responsibility of the authors and does not necessarily represent the official views of the NIH.

### Competing interests

The authors have declared that no competing interests exist.

---

**Fig. S1. 15 mM MG inhibits growth in a variable and strain-dependent manner.** Representative growth kinetics for *C. lusitaniae* strains grown in YPD in the absence **(A, B)** or presence **(C,D)** of 15 mM MG. S18 **(A,C)** or L17 **(B,D)** parental (black) and isogenic *mgd1∆* (magenta), *mgd2∆* (teal), and *mgd1∆*/*mgd2∆* (purple) are shown. One representative experiment out of four **(B,D)** or five **(A,C)** independent experiments is shown. Error bars indicate technical replicates from the same experiment.

**Fig. S2. *MGD1*, *MGD2*, and *MRR1* play a role in MG catabolism.** *C. lusitaniae* S18 wild type (WT), were grown a microtiter dish at 37C for 36 hours in YNB medium in supplemented with either 5 mM glucose **(A)** or 5 mM MG **(B)**, then final OD_600_ was measured. One-way ANOVA was used for statistical evaluation; *, p < 0.05. Data shown represent the mean final OD_600_ from each of five independent experiments. Connecting lines indicate each experiment.

**Fig. S3. *MRR1* and *CAP1* play a role in MG-dependent *MDR1* induction and FLZ resistance in *C. lusitaniae* isolate L17. (A**) Induction of *MDR1* in L17 WT (black), *mrr1Δ* (orange), and *cap1Δ* (green) following 15 minutes of exposure to 5 mM MG in YPD-grown exponential phase cells. Data shown represent the mean ± SD from three independent experiments. **(B)** Growth curves for L17 WT in YPD alone (blue), or with 5 mM MG (red), FLZ (green), or FLZ + 5 mM MG (light purple). **(C)** Final OD_600_ at 16 hours of growth for indicated strains in YPD alone or with 5 mM MG and/or FLZ. Data shown represent the mean ± SD from three independent experiments. Three-way ANOVA was used for statistical evaluation in **(A)** and **(C)**.

**Fig. S4. Loss of *CAP1* increased sensitivity to high concentrations of exogenous MG regardless of whether *MRR1* was present.** *C. lusitaniae* S18 (black), *cap1∆* (green), and *mrr1∆*/*cap1∆* (yellow) were grown in YPD alone **(A**) or with 15 mM MG **(B)**. One representative experiment out of three is shown.

## References

1. Lamoth F, Lockhart SR, Berkow EL, Calandra T. Changes in the epidemiological landscape of invasive candidiasis. J Antimicrob Chemother. 2018;73(suppl_1):i4–i13.

2. Arendrup MC. Epidemiology of invasive candidiasis. Curr Opin Crit Care. 2010;16(5):445–52.

3. Nucci M, Perfect JR. When primary antifungal therapy fails. Clin Infect Dis. 2008;46(9):1426–33.

4. Hiller D, Sanglard D, Morschhauser J. Overexpression of the *MDR1* gene is sufficient to confer increased resistance to toxic compounds in *Candida albicans*. Antimicrob Agents Chemother. 2006;50(4):1365–71.

5. Jin L, Cao Z, Wang Q, Wang Y, Wang X, Chen H, et al. *MDR1* overexpression combined with *ERG11* mutations induce high-level fluconazole resistance in *Candida tropicalis* clinical isolates. BMC Infect Dis. 2018;18(1):162.

6. Wirsching S, Moran GP, Sullivan DJ, Coleman DC, Morschhauser J. *MDR1*-mediated drug resistance in *Candida dubliniensis*. Antimicrob Agents Chemother. 2001;45(12):3416–21.

7. Demers EG, Biermann AR, Masonjones S, Crocker AW, Ashare A, Stajich JE, et al. Evolution of drug resistance in an antifungal-naive chronic *Candida lusitaniae* infection. Proc Natl Acad Sci U S A. 2018;115(47):12040–5.

8. Dunkel N, Blass J, Rogers PD, Morschhauser J. Mutations in the multi-drug resistance regulator *MRR1*, followed by loss of heterozygosity, are the main cause of *MDR1* overexpression in fluconazole-resistant *Candida albicans* strains. Mol Microbiol. 2008;69(4):827–40.

9. Schubert S, Rogers PD, Morschhauser J. Gain-of-function mutations in the transcription factor *MRR1* are responsible for overexpression of the *MDR1* efflux pump in fluconazole-resistant *Candida dubliniensis* strains. Antimicrob Agents Chemother. 2008;52(12):4274–80.

10. Morschhauser J, Barker KS, Liu TT, Bla BWJ, Homayouni R, Rogers PD. The transcription factor Mrr1p controls expression of the *MDR1* efflux pump and mediates multidrug resistance in *Candida albicans*. PLoS Pathog. 2007;3(11):e164.

11. Liu Z, Myers LC. *Candida albicans* Swi/Snf and Mediator complexes differentially regulate Mrr1-induced *MDR1* expression and fluconazole resistance. Antimicrob Agents Chemother. 2017;61(11).

12. Mogavero S, Tavanti A, Senesi S, Rogers PD, Morschhauser J. Differential requirement of the transcription factor Mcm1 for activation of the *Candida albicans* multidrug efflux pump *MDR1* by its regulators Mrr1 and Cap1. Antimicrob Agents Chemother. 2011;55(5):2061–6.

13. Schubert S, Barker KS, Znaidi S, Schneider S, Dierolf F, Dunkel N, et al. Regulation of efflux pump expression and drug resistance by the transcription factors Mrr1, Upc2, and Cap1 in *Candida albicans*. Antimicrob Agents Chemother. 2011;55(5):2212–23.

14. Karababa M, Coste AT, Rognon B, Bille J, Sanglard D. Comparison of gene expression profiles of *Candida albicans* azole-resistant clinical isolates and laboratory strains exposed to drugs inducing multidrug transporters. Antimicrob Agents Chemother. 2004;48(8):3064–79.

15. Hoehamer CF, Cummings ED, Hilliard GM, Morschhauser J, Rogers PD. Proteomic analysis of Mrr1p- and Tac1p-associated differential protein expression in azole-resistant clinical isolates of *Candida albicans*. Proteomics Clin Appl. 2009;3(8):968–78.

16. Rogers PD, Barker KS. Genome-wide expression profile analysis reveals coordinately regulated genes associated with stepwise acquisition of azole resistance in *Candida albicans* clinical isolates. Antimicrob Agents Chemother. 2003;47(4):1220–7.

17. Silva AP, Miranda IM, Guida A, Synnott J, Rocha R, Silva R, et al. Transcriptional profiling of azole-resistant *Candida parapsilosis* strains. Antimicrob Agents Chemother. 2011;55(7):3546–56.

18. Kannan A, Asner SA, Trachsel E, Kelly S, Parker J, Sanglard D. Comparative genomics for the elucidation of multidrug resistance in *Candida lusitaniae*. mBio. 2019;10(6).

19. Zuin A, Vivancos AP, Sanso M, Takatsume Y, Ayte J, Inoue Y, et al. The glycolytic metabolite methylglyoxal activates Pap1 and Sty1 stress responses in *Schizosaccharomyces pombe*. The Journal of biological chemistry. 2005;280(44):36708–13.

20. Takatsume Y, Izawa S, Inoue Y. Methylglyoxal as a signal initiator for activation of the stress-activated protein kinase cascade in the fission yeast *Schizosaccharomyces pombe*. The Journal of biological chemistry. 2006;281(14):9086–92.

21. Lu J, Randell E, Han Y, Adeli K, Krahn J, Meng QH. Increased plasma methylglyoxal level, inflammation, and vascular endothelial dysfunction in diabetic nephropathy. Clinical biochemistry. 2011;44(4):307–11.

22. Wang XJ, Ma SB, Liu ZF, Li H, Gao WY. Elevated levels of alpha-dicarbonyl compounds in the plasma of type II diabetics and their relevance with diabetic nephropathy. J Chromatogr B Analyt Technol Biomed Life Sci. 2019;1106-1107:19–25.

23. McLellan AC, Thornalley PJ, Benn J, Sonksen PH. Glyoxalase system in clinical diabetes mellitus and correlation with diabetic complications. Clin Sci (Lond). 1994;87(1):21–9.

24. Brenner T, Fleming T, Uhle F, Silaff S, Schmitt F, Salgado E, et al. Methylglyoxal as a new biomarker in patients with septic shock: an observational clinical study. Crit Care. 2014;18(6):683.

25. Mukhopadhyay S, Ghosh A, Kar M. Methylglyoxal increase in uremia with special reference to snakebite-mediated acute renal failure. Clin Chim Acta. 2008;391(1-2):13–7.

26. Lapolla A, Flamini R, Lupo A, Arico NC, Rugiu C, Reitano R, et al. Evaluation of glyoxal and methylglyoxal levels in uremic patients under peritoneal dialysis. Ann N Y Acad Sci. 2005;1043:217–24.

27. Karg E, Papp F, Tassi N, Janaky T, Wittmann G, Turi S. Enhanced methylglyoxal formation in the erythrocytes of hemodialyzed patients. Metabolism. 2009;58(7):976–82.

28. Odani H, Shinzato T, Usami J, Matsumoto Y, Brinkmann Frye E, Baynes JW, et al. Imidazolium crosslinks derived from reaction of lysine with glyoxal and methylglyoxal are increased in serum proteins of uremic patients: evidence for increased oxidative stress in uremia. FEBS Lett. 1998;427(3):381–5.

29. Zhang MM, Ong CL, Walker MJ, McEwan AG. Defence against methylglyoxal in Group A *Streptococcus*: a role for Glyoxylase I in bacterial virulence and survival in neutrophils? Pathog Dis. 2016;74(2).

30. Stewart BJ, Navid A, Kulp KS, Knaack JLS, Bench G. D-Lactate production as a function of glucose metabolism in *Saccharomyces cerevisiae*. Yeast. 2013;30(2):81–91.

31. Martins AM, Cordeiro CA, Ponces Freire AM. In situ analysis of methylglyoxal metabolism in *Saccharomyces cerevisiae*. FEBS letters. 2001;499(1-2):41–4.

32. Thornalley PJ. The glyoxalase system: new developments towards functional characterization of a metabolic pathway fundamental to biological life. The Biochemical journal. 1990;269(1):1–11.

33. Ray M, Ray S. Purification and partial characterization of a methylglyoxal reductase from goat liver. Biochimica et biophysica acta. 1984;802(1):119–27.

34. Maeta K, Izawa S, Okazaki S, Kuge S, Inoue Y. Activity of the Yap1 transcription factor in *Saccharomyces cerevisiae* is modulated by methylglyoxal, a metabolite derived from glycolysis. Mol Cell Biol. 2004;24(19):8753–64.

35. Inoue Y, Kimura A. Identification of the structural gene for glyoxalase I from *Saccharomyces cerevisiae*. J Biol Chem. 1996;271(42):25958–65.

36. Chen CN, Porubleva L, Shearer G, Svrakic M, Holden LG, Dover JL, et al. Associating protein activities with their genes: rapid identification of a gene encoding a methylglyoxal reductase in the yeast *Saccharomyces cerevisiae*. Yeast. 2003;20(6):545–54.

37. Kwak MK, Ku M, Kang SO. Inducible NAD(H)-linked methylglyoxal oxidoreductase regulates cellular methylglyoxal and pyruvate through enhanced activities of alcohol dehydrogenase and methylglyoxal-oxidizing enzymes in glutathione-depleted *Candida albicans*. Biochimica et biophysica acta General subjects. 2018;1862(1):18–39.

38. Stajich JE, Harris T, Brunk BP, Brestelli J, Fischer S, Harb OS, et al. FungiDB: an integrated functional genomics database for fungi. Nucleic Acids Res. 2012;40(Database issue):D675–81.

39. Basenko EY, Pulman JA, Shanmugasundram A, Harb OS, Crouch K, Starns D, et al. FungiDB: An Integrated Bioinformatic Resource for Fungi and Oomycetes. J Fungi (Basel). 2018;4(1).

40. Murata K, Inoue Y, Saikusa T, Watanabe K, Fukuda Y, Shimosaka M, et al. Metabolism of alpha-ketoaldehydes in yeasts - inducible formation of methylglyoxal reductase and its relation to growth arrest of *Saccharomyces cerevisiae*. J Ferment Bioeng. 1986;64(1):1–4.

41. Znaidi S, Barker KS, Weber S, Alarco AM, Liu TT, Boucher G, et al. Identification of the *Candida albicans* Cap1p regulon. Eukaryot Cell. 2009;8(6):806–20.

42. Thornalley PJ. Pharmacology of methylglyoxal: formation, modification of proteins and nucleic acids, and enzymatic detoxification--a role in pathogenesis and antiproliferative chemotherapy. Gen Pharmacol. 1996;27(4):565–73.

43. Penninckx MJ, Jaspers CJ, Legrain MJ. The glutathione-dependent glyoxalase pathway in the yeast *Saccharomyces cerevisiae*. J Biol Chem. 1983;258(10):6030–6.

44. Harry JB, Oliver BG, Song JL, Silver PM, Little JT, Choiniere J, et al. Drug-induced regulation of the *MDR1* promoter in *Candida albicans*. Antimicrob Agents Chemother. 2005;49(7):2785–92.

45. Calabrese D, Bille J, Sanglard D. A novel multidrug efflux transporter gene of the major facilitator superfamily from *Candida albicans* (*FLU1*) conferring resistance to fluconazole. Microbiology. 2000;146 (Pt 11):2743–54.

46. Li R, Kumar R, Tati S, Puri S, Edgerton M. *Candida albicans* Flu1-mediated efflux of salivary histatin 5 reduces its cytosolic concentration and fungicidal activity. Antimicrob Agents Chemother. 2013;57(4):1832–9.

47. Rodrigues CF, Rodrigues ME, Henriques M. *Candida* sp. infections in patients with diabetes mellitus. J Clin Med. 2019;8(1).

48. Pyrgos V, Ratanavanich K, Donegan N, Veis J, Walsh TJ, Shoham S. *Candida* bloodstream infections in hemodialysis recipients. Med Mycol. 2009;47(5):463–7.

49. Jawale C, Ramani K, Biswas PS. Defect in neutrophil function accounts for impaired anti-fungal immunity in kidney dysfunction. J Immunol. 2018;200(1).

50. Beisswenger PJ, Howell SK, Touchette AD, Lal S, Szwergold BS. Metformin reduces systemic methylglyoxal levels in type 2 diabetes. Diabetes. 1999;48(1):198–202.

51. Ogasawara Y, Tanaka R, Koike S, Horiuchi Y, Miyashita M, Arai M. Determination of methylglyoxal in human blood plasma using fluorescence high performance liquid chromatography after derivatization with 1,2-diamino-4,5-methylenedioxybenzene. J Chromatogr B Analyt Technol Biomed Life Sci. 2016;1029-1030:102–5.

52. Vander Jagt DL, Hunsaker LA. Methylglyoxal metabolism and diabetic complications: roles of aldose reductase, glyoxalase-I, betaine aldehyde dehydrogenase and 2-oxoaldehyde dehydrogenase. Chem Biol Interact. 2003;143-144:341–51.

53. Liu J, Wang R, Desai K, Wu L. Upregulation of aldolase B and overproduction of methylglyoxal in vascular tissues from rats with metabolic syndrome. Cardiovasc Res. 2011;92(3):494–503.

54. Masterjohn C, Park Y, Lee J, Noh SK, Koo SI, Bruno RS. Dietary fructose feeding increases adipose methylglyoxal accumulation in rats in association with low expression and activity of glyoxalase-2. Nutrients. 2013;5(8):3311–28.

55. Alarco AM, Raymond M. The bZip transcription factor Cap1p is involved in multidrug resistance and oxidative stress response in *Candida albicans*. J Bacteriol. 1999;181(3):700–8.

56. Znaidi S, Weber S, Al-Abdin OZ, Bomme P, Saidane S, Drouin S, et al. Genomewide location analysis of *Candida albicans* Upc2p, a regulator of sterol metabolism and azole drug resistance. Eukaryot Cell. 2008;7(5):836–47.

57. Kettle AJ, Turner R, Gangell CL, Harwood DT, Khalilova IS, Chapman AL, et al. Oxidation contributes to low glutathione in the airways of children with cystic fibrosis. The European respiratory journal. 2014;44(1):122–9.

58. Dickerhof N, Pearson JF, Hoskin TS, Berry LJ, Turner R, Sly PD, et al. Oxidative stress in early cystic fibrosis lung disease is exacerbated by airway glutathione deficiency. Free radical biology & medicine. 2017;113:236–43.

59. Yamauchi S, Kiyosawa N, Ando Y, Watanabe K, Niino N, Ito K, et al. Hepatic transcriptome and proteome responses against diethyl maleate-induced glutathione depletion in the rat. Arch Toxicol. 2011;85(9):1045–56.

60. Urban N, Tsitsipatis D, Hausig F, Kreuzer K, Erler K, Stein V, et al. Non-linear impact of glutathione depletion on *C. elegans* life span and stress resistance. Redox Biol. 2017;11:502–15.

61. Enkvetchakul B, Bottje WG. Influence of diethyl maleate and cysteine on tissue glutathione and growth in broiler chickens. Poult Sci. 1995;74(5):864–73.

62. Mitchell JB, Russo A, Biaglow JE, Mcpherson SJ. Cellular glutathione depletion by diethyl maleate or buthionine sulfoximine and its effects on the oxygen enhancement ratio. Radiat Res. 1983;94(3):612-.

63. Zheng J, Hu CL, Shanley KL, Bizzozero OA. Mechanism of protein carbonylation in glutathione-depleted rat brain slices. Neurochem Res. 2018;43(3):609–18.

64. Yadav SK, Singla-Pareek SL, Ray M, Reddy MK, Sopory SK. Methylglyoxal levels in plants under salinity stress are dependent on glyoxalase I and glutathione. Biochem Biophys Res Commun. 2005;337(1):61–7.

65. Shen XX, Zhou XF, Kominek J, Kurtzman CP, Hittinger CT, Rokas A. Reconstructing the backbone of the *Saccharomycotina* yeast phylogeny using genome-scale data. G3-Genes Genom Genet. 2016;6(12):3927–39.

66. Pathirana RU, Friedman J, Norris HL, Salvatori O, McCall AD, Kay J, et al. Fluconazole-resistant *Candida auris* Is susceptible to salivary histatin 5 killing and to intrinsic host defenses. Antimicrob Agents Ch. 2018;62(2).

67. Grahl N, Demers EG, Crocker AW, Hogan DA. Use of RNA-protein complexes for genome editing in non-albicans *Candida* species. mSphere. 2017;2(3).

